# Environmental factors and microbe-microbe interactions drive the structure of the core microbiota of terrestrial microalgae

**DOI:** 10.64898/2026.09.06.749702

**Authors:** Vincent Garrigues, Rui Guan, Nathan Preteseille, Iréne Leccia, Edwin Wagner, Fabrice Roux, Paloma Durán

**Affiliations:** LIPME, Centre National de la Recherche Scientifique, Institut National de la Recherche Agronomique, Castanet-Tolosan, France; Quadram Institute Bioscience, Norwich Research Park, Norwich, England, UK

**Keywords:** core microbiota, plant microbiota, environmental drivers, microbial networks, field survey

## Abstract

Plants and other photosynthetic organisms interact with their environment and surrounding microbiota through specialized associations. A global core microbiota has been proposed at high taxonomic levels, such as the order level. However, it remains unclear which environmental factors and how microbe-microbe interactions drive variation of this core microbiota at lower taxonomic resolution. Here, we leveraged the environmental diversity of 141 sites across the southwest of France to characterize algal populations, and their associated bacterial and fungal microbiota. We then performed a meta-analysis, combining these data with published datasets to formally identify the global core microbiota of terrestrial photosynthetic organisms, which comprises seven bacterial and five fungal orders. We next investigated diversity within this core microbiota and the environmental drivers shaping site-specific community composition. While environmental factors have a low impact on the total relative abundance of core orders, the core microbiota at the ASV-level is impacted by climatic factors, edaphic factors, and plant community descriptors. Using interaction network analysis, we finally explored how microbe-microbe interactions contribute to the assembly of stable core communities. Our results show that core ASVs occupy central positions in algal-associated microbial networks and that distinct core orders drive site-specific variation in core microbiota structure.

Together, these findings highlight the importance of both environmental context and microbial interactions in shaping the composition and stability of the core microbiota associated with photosynthetic organisms.

## Introduction

In natural environments, microbial communities are found in association with eukaryotic hosts. These interactions affect host performance under natural conditions and may be influenced by environmental conditions ^[1, 2]^. For plant rhizosphere-associated communities, environmental factors such as soil pH, soil water content, or plant community composition have been identified as important drivers of microbial community structure ^[3, 4]^. At the same time, it has been shown that a high-taxonomic-level subset of these rhizospheric microbial communities is systematically found associated with all photosynthetic organisms, regardless of the species or the habitat in which they are found, the so-called core microbiota ^[5, 6, 7]^. The fact that these microbial members are present across multiple host plants and environments suggests that they are well adapted to interacting with photosynthetic organisms and with one another. Furthermore, their consistent presence suggests a beneficial impact on their host. While the potential effects of a global core microbiome on host performance remain largely underexplored, several reports have described host- or treatment-specific consensus microbiota sets with beneficial effects on their hosts. For example, by studying multiple grass species grown in an experimental field, a set of 278 taxa was identified, which correlated with an increase in nitrogen-cycling genes in the rhizosphere ^[8]^. In Qiu *et al.*,^[9]^, the authors identified 33 bacterial OTUs as common microbial members of the cotton rhizosphere that positively correlated with the health status of their host. Recently, it has been shown that 25 bacterial strains are specifically enriched in *Hyaloperonospora*-infected plants, thereby suggesting a plant protection against the pathogen ^[10, 11]^. Therefore, it is tempting to hypothesize that a global core microbiota of all photosynthetic organisms also has important roles for plant performance and/or environmental adaptation.

However, it remains unclear whether environmental factors or microbe-microbe interactions drive variation of this core microbiota, in particular at lower taxonomic resolution. There are many examples where the rhizosphere microbiota was studied across multiple habitats and where the diversity of the core microbiota could be studied (e.g. in ^[3, 12]^). However, the variability of environmental factors in these surveys is limited and it remains costly, labor-intensive and disturbing to the natural environment to perform more comprehensive field surveys. In addition, our understanding of the interactions of core microbiota members between them, with other members of the local microbiota, and the environment remains limited. Advances in high-throughput sequencing have generated vast datasets from many terrestrial environments, which allows exploring the possible diversity within very conserved core microbiota groups, but also enables the growing application of graph theory in microbial ecology. Network analysis has emerged as a powerful tool to uncover complex associations within microbial communities, and how environmental factors can affect network properties, thereby revealing interactions between the aforementioned core microbiota, and microecosystem-level dynamics that are often obscured in traditional analyses ^[13]^.

In previous work, it was demonstrated that *Chlamydomonas reinhardtii* and other subaerial unicellular algae, also interact with the core microbiota of land plants, suggesting that these interactions are conserved across photosynthetic organisms ^[6]^. Therefore, using subaerial algae as focal host to understand the core microbiota represents an interesting opportunity, since it allows fast sampling, with very little disturbance to the natural environment. Surprisingly, despite these advantages, and in contrast to marine algal communities, which have been broadly described over the last decade ^[14, 15]^, subaerial algal communities are very poorly represented in the literature ^[16, 17]^.

Here, we performed a survey across soils of 141 natural sites across the southwest of France previously characterized for a large set of abiotic (climate and edaphic properties) and biotic (plant communities) factors ^[18, 19, 20]^. We described terrestrial algal populations using 18S amplicon sequencing and profiled bacterial and fungal communities associated with these algal communities by 16S and ITS amplicon sequencing, respectively. We take these and publicly available datasets to define a global core microbiota of photosynthetic organisms, shared from terrestrial microalgae to land plants. Leveraging these datasets, we asked: 1) How prevalent are subaerial algae across a diversity of natural environments and which environmental factors shape their natural distribution? 2) Can we find the core microbiota of photosynthetic organisms in association with subaerial algal populations? If so, which environmental factors determine the core microbiota structure and its potential variation at the ASV-level? 3) Are microbe-microbe interactions important drivers of the structure of the core microbiota in natural sites?

## Results

### Algal populations are prevalent across natural sites but depend on environmental variation

To estimate whether algal populations are common in natural environments, we selected 141 natural sites across the southwest of France with a diverse soil and plant community composition ^[20]^. Additionally, we retrieved meteorological data from nearby weather stations from four weeks before harvesting dates (**Supplementary Table 1, Supplementary Figure 1**). As previously shown, soil native algae communities are distinct between the soil surface and deeper soil layers ^[6]^ (**Supplementary Figure 2a**), and are generally more abundant in the photosynthetic fraction of the soil, i.e. on the soil surface, which may be more accentuated in some natural soils compared to others (**Supplementary Figure 2b**). We sampled four surface soil samples per site, where obvious green non-moss patches were visible. Amplicon sequencing of the 18S rRNA marker gene showed that green algae (Chlorophyta) are among the four most abundant eukaryotic groups found across all sites, together with Phragmoplastophyta (plant DNA), Cercozoa (protists) and Ascomycota (fungi) (**Supplementary Figure 3**). Among green algae, the most prevalent algal class is Chlorophyceae, followed by Trebouxiophyceae, Klebsormidiophyceae and Ulvophyceae, which are well distributed across all sites harvested (**Figure 1a**). Interestingly, members of the *Chlamydomonas* genus have been studied for decades under laboratory conditions, but there is limited knowledge on their distribution in natural conditions. In this survey, members of *Chlamydomonas* species were found at 66 sites, including *C. reinhardtii*, *C. moewusii* and *C. bacca* (**Supplementary Figure 4**).

**Figure 1:**
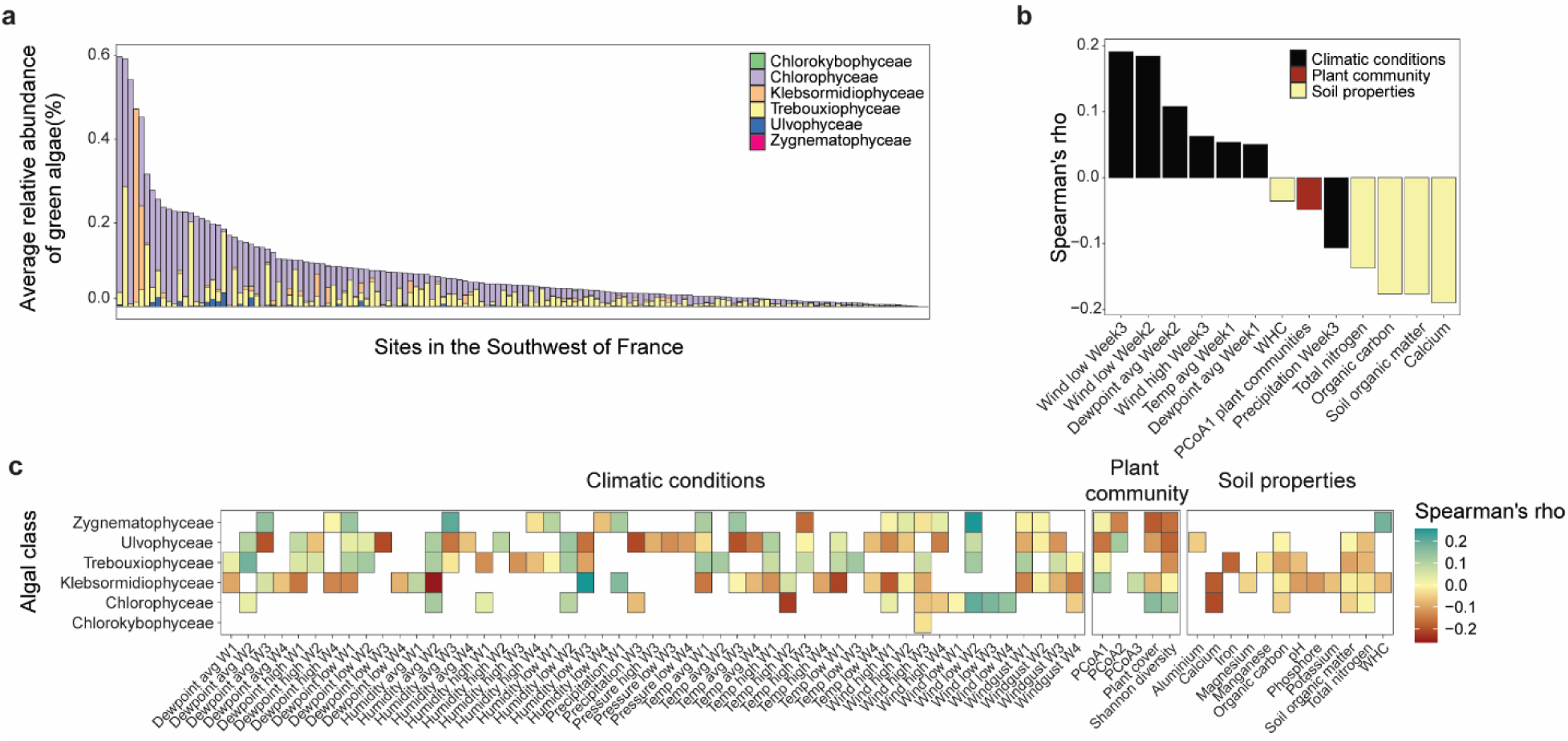
Subaerial algae are abundant across multiple sites of the southwest of France. **a)** Average relative abundance of green algae across all surveyed sites, color coded by their taxonomic affiliation at class level (n=4, for each site). **b)** Correlation between total green algae abundance and environmental factors. Only significant correlations are shown here (Spearman’s correlation, *P* < 0.05, FDR-corrected), color coded by the type of environmental factor (climatic conditions, plant community or soil properties). **c)** Correlation between the relative abundance of algal classes and environmental factors. Only significant correlations are shown here (Spearman’s correlation, *P* < 0.05, FDR-corrected), color-coded based on the strength of the correlation.

Previous studies have shown that climatic conditions, soil characteristics or plant cover are important drivers of algal biomass in natural conditions ^[17]^. We made use of the available environmental variables for the surveyed sites (**Supplementary Table 1**), which highlighted that specific soil properties, such as organic carbon, calcium or nitrogen negatively correlate with total algal abundance (**Figure 1b**). Similarly, precipitation and plant community predictors also negatively impacted algal abundance. On the other hand, climatic conditions such as wind, temperature and dew point positively correlate with the abundance of native algal populations (**Figure 1b**). Interestingly, not all algal classes are equally affected by environmental factors (**Figure 1c**). Chlorophyceae are relatively stable across sites and associated with a handful of factors, although strongly driven by wind, plant communities, and calcium. On the other hand, Klebsormidiophyceae, Trebouxiophyceae and Ulvophyceae are associated with multiple environmental factors, such as dew point, humidity, pressure, temperature, wind, plant communities, organic carbon, and total nitrogen (**Figure 1c**). Because members of the Chlorokybophyceae and Zygnematophyceae families were found only in three of the surveyed sites, we lacked statistical power to estimate correlations with environmental factors. Finally, we grouped our sites into environmental clusters (ECs, **Methods**) to assess whether algal communities are driven by general ecological profiles. Environmental clusters were defined based on similarities of climatic conditions, edaphic factors and plant descriptors (**Methods**, **Supplementary Figure 1**). PERMANOVA analysis showed that algal communities’ structure is not driven by the three identified ECs, suggesting that algal communities are predominantly structured by a combination of site-specific environmental factors (**Supplementary Figure 5**, *P* = 0.97).

### Environmental factors affect the ASV-level structure of the core microbiota of algal populations

The core microbiota is a concept largely used in most host-associated microbiota research fields, and corresponds to the fraction of the total microbiota shared across hosts, environments, and/or over time ^[21]^. Importantly, this core microbiota may vary between studies, because each study relies on different statistical thresholds, taxonomic levels, and may contain more or less host species and natural sites ^[22]^. To statistically and robustly define the core microbiota of terrestrial photosynthetic organisms, we first retrieved published datasets from roots and phycospheres of 46 photosynthetic species from 54 habitats (**Supplementary Table 2**). Additionally, we performed 16S rRNA and ITS amplicon sequencing on the same samples described above to describe the bacterial and fungal communities associated with wild algal populations from the southwest of France. Thereby, we incremented the dataset to 46 individual photosynthetic species, 141 terrestrial algal populations and 195 habitats (**Methods**). Relative abundances of bacterial and fungal members were aggregated at the order level, and we applied the statistical framework described by Shade & Stopnisek ^[23]^, to identify microbial taxa which are prevalent, abundant, and have a significant impact on microbial community structure. Thereby, the core microbiota was defined as those orders that were found in all hosts and sites (100% occupancy), with a relative abundance >1% and that had a significant impact on the overall microbial community structure (an increase in the community Bray-Curtis dissimilarity of equal or greater than 2%) (**Methods**). We identified seven bacterial (Bulkholderiales, Caulobacterales, Microtrichales, Propionibacteriales, Rhizobiales, Solirubrobacterales, Sphingomonadales) and five fungal orders (Chaetothyriales, Helotiales, Hypocreales, Pleosporales, Xylariales) as members of the core microbiota of photosynthetic organisms (**Figure 2**). Among them, several of those were already described in previous work, namely Bulkholderiales, Caulobacterales, Rhizobiales and Sphingomonadales (e.g. in ^7, 24^).

**Figure 2:**
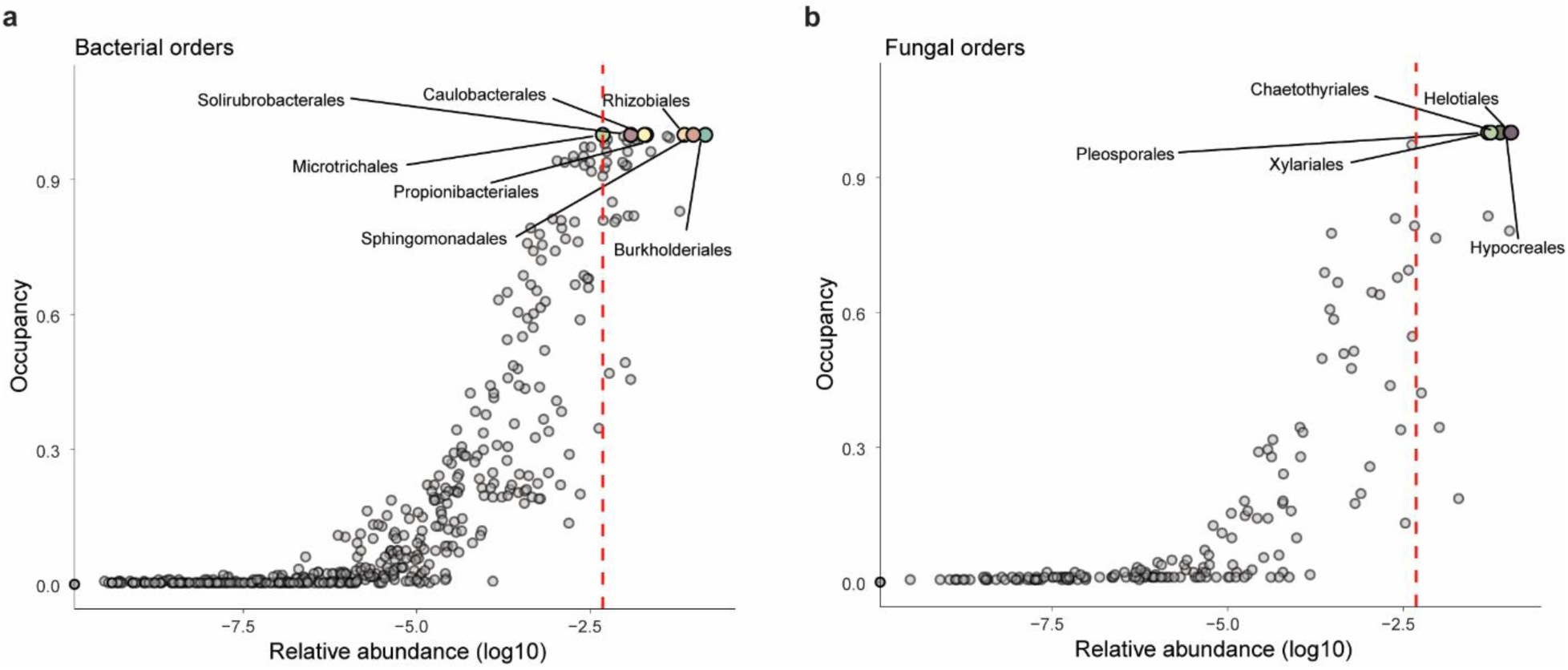
The core microbiota of photosynthetic organisms. Abundance-occupancy plots for bacterial (**a**) and fungal (**b**) orders. Those orders which are abundantly present in all samples and impact microbial community structure (elbow method, dashed red line) are considered as core orders and are highlighted here.

To identify the potential environmental drivers of these core groups, we focused on our algal-associated dataset. We observed that the relative abundances of core orders depends on the site (**Figure 3a, c**). While bacterial core orders are stably found between 21.5% to 35.2% of the total bacterial relative abundance, fungal core orders range from 7.9% to 48.3% of the total fungal relative abundance (5th–95th percentiles), suggesting that among core members, bacteria are more stable across environments than fungi (**Figure 3a, c**). Both bacterial and fungal communities were different between algal-associated soil surfaces and deeper soil layers (PERMANOVA, 26% and 28% of the total variance explained by site and soil layer, respectively, *P* < 0.001, **Supplementary Figure 2c, e**). When exploring specifically the relative abundance of individual core orders, only three bacterial orders (Burkholderiales, Caulobacteriales and Sphingomonadales) and one fungal order (Xylariales) were significantly more abundant in algal-associated soil layers (**Supplementary Figure 2d, f**, *P* < 0.05, Wilcoxon Test, Holm-corrected).

**Figure 3:**
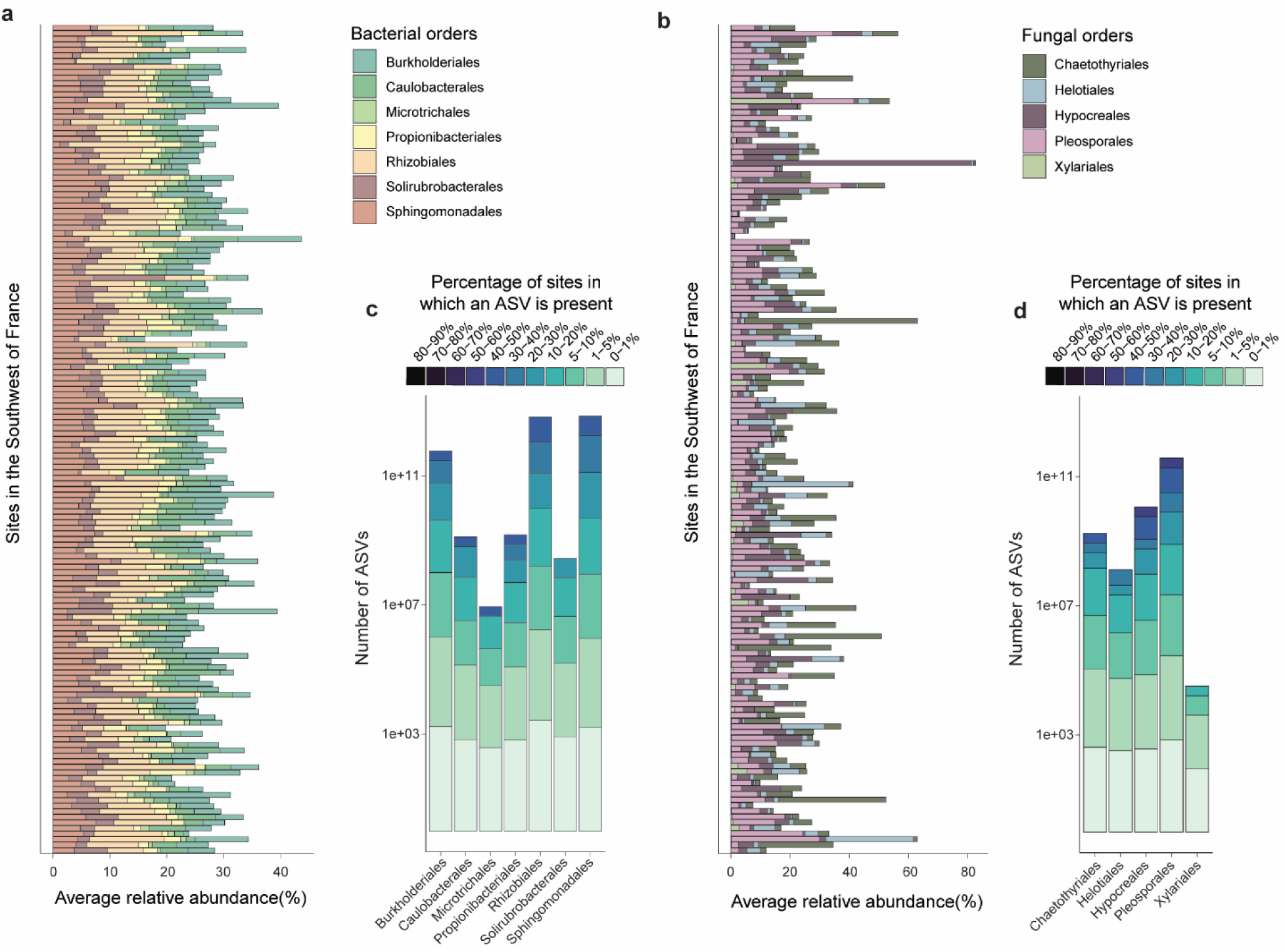
The core microbiota of photosynthetic organisms associates with wild algal populations. Average relative abundance of bacterial **(a**) and fungal **(c)** core orders across 141 sites (n=4, for each site). **b, d)** Number of core microbiota ASVs found across all the surveyed sites, color-coded by percentage of sites in which they are found. Due to the high number of ASVs found in a single site, values have been log10-transformed.

We next inspected lower taxonomic levels (i.e., at ASV level) and observed that not a single ASV was found across all sites surveyed in this study, suggesting that core members are conserved at high taxonomic levels, but that they vary depending on their site at lower taxonomic resolution (**Figure 3b, d**). The diversity of surveyed environments allowed us to assess which factors may be important for core microbiota ASV-level structure (**Supplementary Figure 1**). Interestingly, environmental factors have a low impact on the total relative abundance of core orders, but the number of ASVs found per site are highly impacted (**Figure 4**). For example, Burkholderiales and Rhizobiales are among the most abundant core groups across sites, and yet their relative abundance is impacted by very few factors, such as manganese or atmospheric pressure (**Figure 4a, bottom panel**). On the contrary, the number of ASVs of these core groups is driven by multiple environmental factors (**Figure 4a, top panel**). This suggests that the overall relative abundance of these core groups is very resilient to environmental variation, but their potential functions in the community are retained by having a diverse set of ASV members. Indeed, when analyzing the impact of environmental factors on the relative abundance of specific ASVs, we observed that ASVs of each core group are both positively and negatively correlated with different environmental factors (**Figure 4b, Supplementary Figure 6**), which further points towards an ASV-balancing within core groups. Among these factors, low wind one to four weeks before harvesting stood out as it strongly and positively correlates with many core microbiota ASVs.

**Figure 4:**
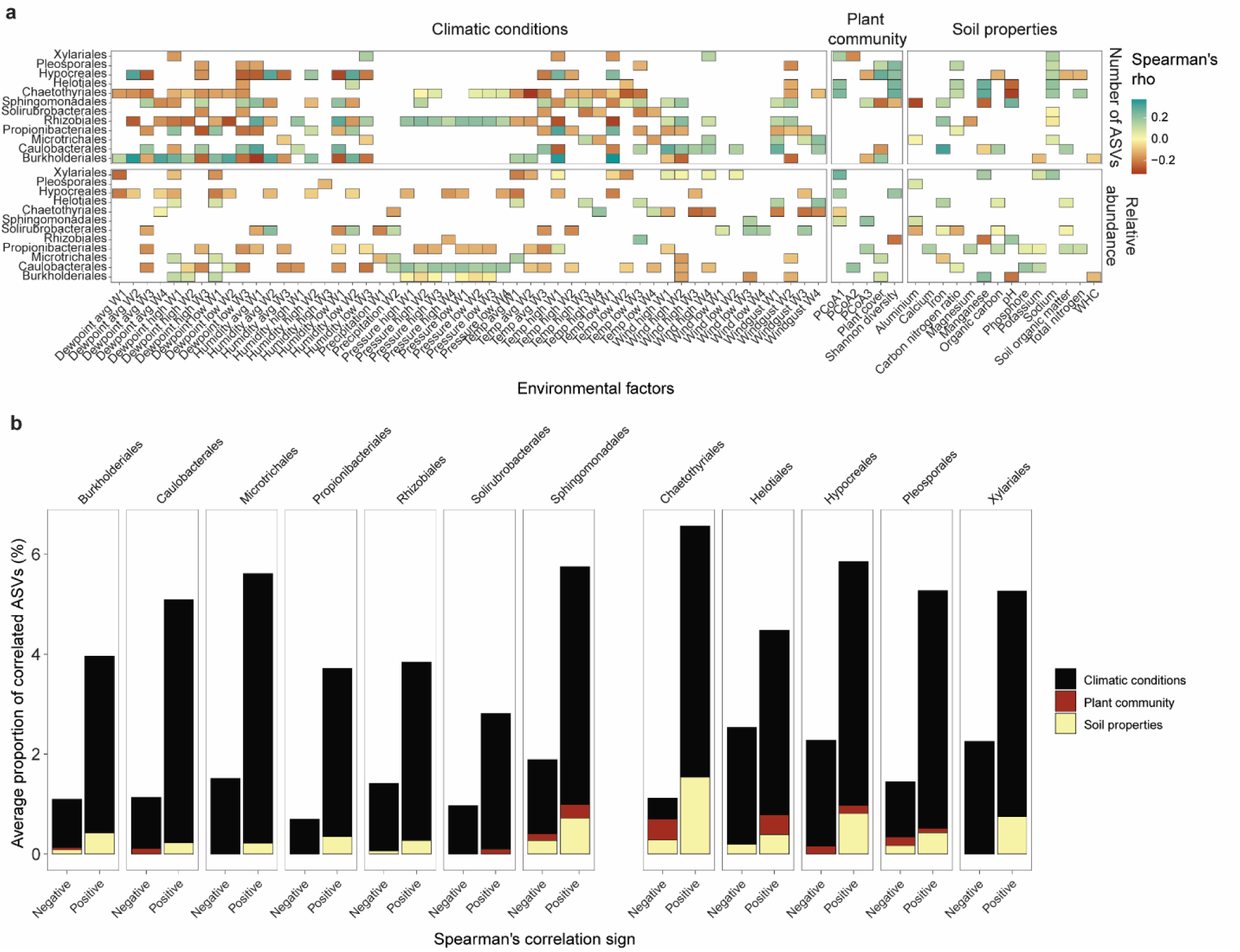
The number of ASVs per core order is affected by environmental factors. **a)** Correlation between environmental factors and the number of ASVs found in a given site (top panel), as well as with the total relative abundance of each core order (bottom panel). Only significant correlations are shown here (Spearman’s correlation, *P* < 0.05, FDR-corrected), color-coded based on the strength of the correlation. **b)** Proportion of ASVs which are significantly correlated (either positive or negative correlation) to environmental factors, out of the total number of ASVs in a given site.

### Network analysis highlights the specific properties of interactions among core members in relationship with environmental variation

Because ASV number is variable and different environmental factors are important for each core group, we decided to study the core microbiota as a complex microbial community, rather than individual members, by constructing networks of interactions. We first built a network by considering all ASVs, and investigated whether core members have different network properties from other microbes. Overall, core nodes have significantly lower closeness centrality than expected by chance (permutation test, *P* < 0.01, FDR-corrected, **Methods**), suggesting that core nodes are more central to the network. At the same time, they have a lower degree centrality than expected by chance (permutation test, *P* < 0.01, FDR-corrected, **Methods**), indicating that they form fewer connections to other network nodes. This suggests that while core nodes are well connected, they rely less on connections to other nodes in the community.

Given that environmental factors impact individual ASV’s relative abundances, we further examined whether those factors impact the interactions between community members and are reflected in the microbial networks. We constructed EC-specific networks with bootstrapping and calculated the distance between them using the R package ‘*mina’* ^[13]^ (**Figure 5a**, **Supplementary Figure 7a**, **Methods**). For each EC-specific networks, we observed that positive interactions are predominant within microbial kingdoms, both for core and non-core nodes, but that interkingdom negative interactions (i.e. between bacteria and fungi) are more prevalent between core nodes than they are among non-core across all ECs (**Figure 5a, c, Supplementary Figure 7a).** PERMANOVA test indicated that 54.98% of between-networks differences can be explained by the EC affiliations (**Figure 5b**). Taken together with the observation that environmental factors affect the “intra-order” diversity of core microbiota (**Figure 4**), we hypothesize that both compositional and interactional dynamics of core microbiota are driven by environmental variation. Interestingly, when further exploring network topology, we observed that core nodes have different network properties from those of non-core nodes in EC1, but not in EC2 and EC3 (**Supplementary Figure 7b**, **Supplementary Table 4**). For example, in EC1 core nodes have lower closeness, and they connect to less connected nodes (i.e. lower neighbor connectivity) (**Supplementary Table 4**).

**Figure 5:**
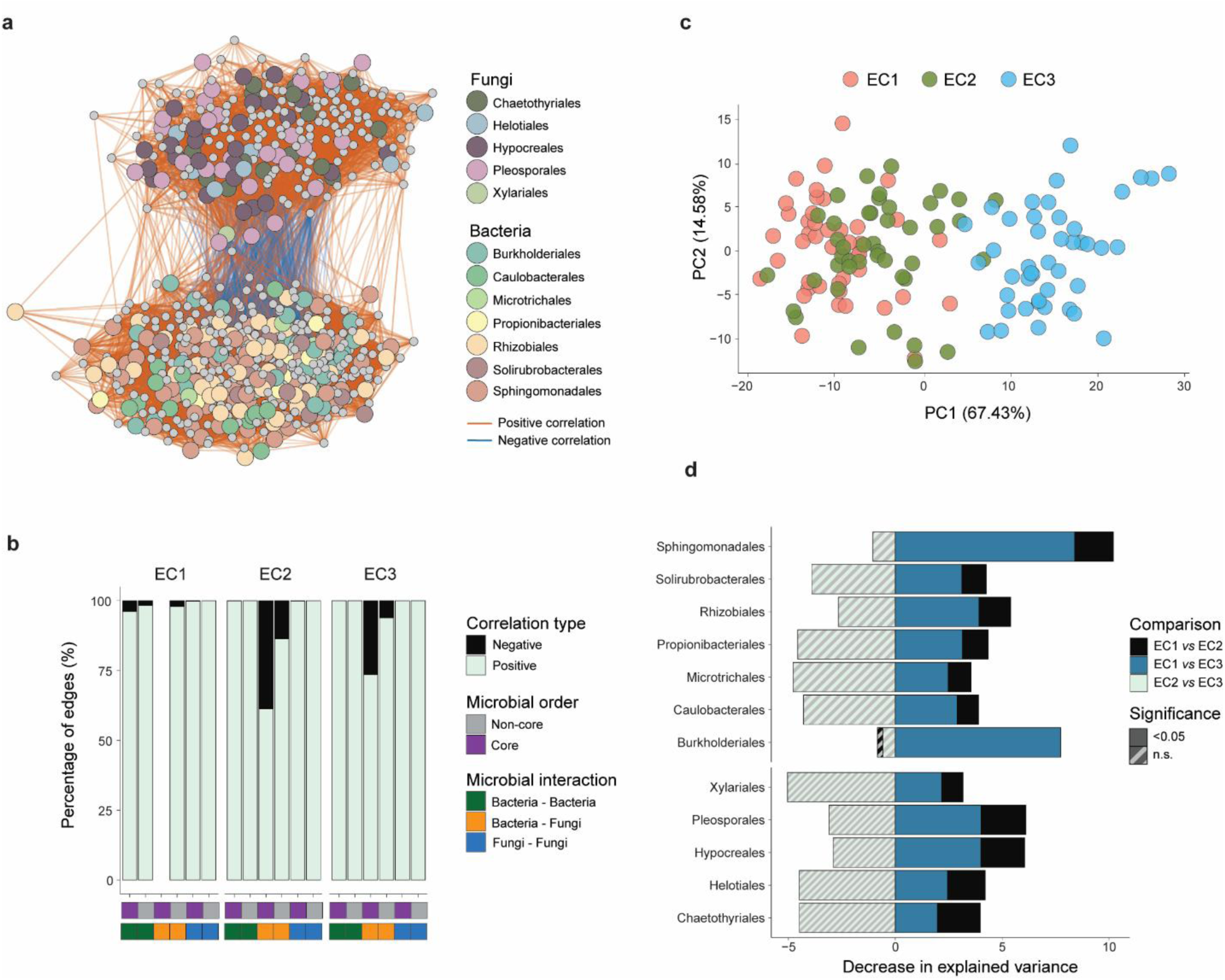
Microbe-microbe interactions structure core microbiota networks. **a)** Network of interactions across natural sites belonging to environmental cluster 1 (EC2 and 3 are shown in **Supplementary** Figure 7a). Colors in nodes indicate whether a given node belongs to a core order (colored), or any other group (grey). Negative correlations are depicted with blue edges, and positive correlations are depicted with orange edges. Only significant correlations are included here (Spearman correlation, *P* < 0.01). **b)** Percentage of edges in each environmental cluster network that have either a negative or positive correlation (only significant correlations are shown, Spearman correlation, *P* < 0.01). Edges are separated between intra-and inter-kingdom interactions, and by core *vs* non-core interactions. **c)** Euclidean distances between network matrices formed by microbiota belonging to the three environmental clusters. Each dot is a bootstrapped network and is color-coded by environmental clusters. **d)** Decrease in variance explained between environmental clusters when removing each core member from the network. Colors indicate significant impact on the variance explained, whereas diagonal stripes indicate non-significant differences.

To identify the core orders that are responsive to the ECs, we further permuted each core order individually to evaluate their contribution to network differentiation (**Methods**). Under the null expectation that microbial occurrence patterns are randomly distributed across ECs, permuting a taxonomic order would not be expected to increase the separation between EC-specific networks. Instead, network distances should remain largely unchanged or fluctuate randomly around their original values. Consistent with this expectation, permutation generally reduced network distances and did not result in any significant increases in network separation (**Figure 5d**). These results indicate that the observed network divergence is maintained by non-random taxonomic association patterns. Interestingly, we observed that the influence of each order on network divergence varies by EC pairs. For example, *Burkholderiales* had no significant effect on the network differentiation, neither between EC1 and EC2 nor between EC2 and EC3. However, the distance between EC1 and EC3 was significantly reduced by 7.74 when removing *Burkholderiales* by permutation, while the mean distance between EC1 and EC3 networks was 23.04 (*P* = 4.04 × 10⁻²², **Figure 5d**). This finding suggests that *Burkholderiales* contributes substantially to the distinct network structure observed between these two environmental clusters, while having little influence on the differentiation of other EC pairs. Additionally, we observed general network topology differences between core orders across ECs. As an example, *Rhizobiales* had a higher betweenness in EC3 in comparison to other other ECs, which may suggest different roles for each core order depending on the EC (**Supplementary Figure 7c**, **Supplementary Table 4**).

## Discussion

The core microbiota is a subset of the diverse microbial communities consistently associated with photosynthetic organisms and has been partially identified in previous studies (e.g. in ^5, 6^). This core microbiota is found in the rhizosphere of land plants and in the phycosphere of soil algae, suggesting common assemblage mechanisms across terrestrial photosynthetic organisms. However, it is well established that the structure and composition of microbial communities are impacted by environmental factors and microbe-microbe interactions (e.g. in ^25, 26^). In this study, we used soil algal communities as a study system to explore the core microbiota, which allows rapid and non-disturbing surveying of multiple natural sites. We acknowledge that isolating a host-associated compartment may not be as straight forward for the phycosphere in comparison to a plant-associated compartment, such as the root endosphere or rhizosphere. Nevertheless, it has been previously demonstrated in controlled laboratory experiments that algal-colonized soil surfaces are enriched in algal-associated microbes, in comparison to non-algae colonized soil surface or deep layers of the soil ^[6]^.

In this study, we have underlined the stability of core microbiota members across 141 natural sites, but we have also leveraged the environmental diversity of these sites to identify environmental factors that drive the ASV-level composition of core orders. In addition, we have studied the effects of environmental factors on core community structure with network analysis, highlighting that network interactions among core groups are defined by environmental drivers, as well as microbe-microbe interactions.

Across our surveyed sites, soil microalgae were prevalent and abundant, as previously demonstrated in other surveys ^[16, 17, 27]^ (**Figure 1a**). The total relative abundances of algal populations were strongly driven by specific environmental drivers. In our analysis, we identified factors such as temperature, plant communities, and soil organic carbon content as important drivers of algal abundances, similar to what has been described in previous works ^[17, 28]^. Here, we deepen the analysis by looking at the drivers of specific algal classes, showing that each algal class is differentially impacted by environmental factors. In addition, while we identified a strong stability of Chlorophyceae across sites (i.e. generalist strategy associated with environmental robustness), the abundance of Klebsormidiophyceae, Trebouxiophyceae and Ulvophyceae appears to be highly modulated by a few environmental drivers (i.e. specialist strategy associated with environmental specialization).

As mentioned above, the bacterial core microbiota had been described previously, considering not only land plants, but also other photosynthetic organisms, such as soil algae ^[6, 7, 24]^. The core microbiota can be defined by considering different parameters: presence/absence, abundance, temporality, ecological relevance, and host adaptation ^[21, 22]^. Most of the core microbiota reports have focused on the presence/absence approach, which does not integrate a measure of overall community impact. Therefore, in this study, we have applied an abundance-occupancy method, coupled with quantification of the impact on the microbial community ^[23]^. Thereby, we gathered published data from root and phycosphere bacterial and fungal microbiota surveys from natural sites and, together with our algal survey, we identified seven bacterial orders and five fungal orders found in association with 47 photosynthetic hosts across 195 sites. We then leveraged the environmental diversity of our algal-associated dataset to find general patterns of the core microbiota structure in natural environments. First, we found that while stable at high taxonomic levels, the core microbiota is variable at the ASV-level (**Figure 3b**, **d**, **Figure 4**). A convergent pattern has been previously observed in the plant microbiota ^[13, 29]^, but also in microbial communities fed with different carbon sources ^[30]^. This suggests that photosynthetic organisms generate a specific niche to which core microbiota members are adapted to colonize across environments, but that ASV-level core microbiota composition may be determined by niche preferences, dispersal limitation, and microbial genomic potential ^[29]^.

Exploration of the impact of environmental drivers on core microbiota ASV-level composition highlighted that soil properties such as pH and manganese are important drivers, as previously demonstrated for plant and soil associated microbiota ^[4, 31, 32, 33]^ (**Figure 4**, **Supplementary Figure 6**). Different climatic variations are important for the relative abundance of both core groups and individual core-ASVs (**Figure 4**, **Supplementary Figure 6**). Interestingly, low wind one to four weeks before harvesting strongly correlates with many core microbiota ASVs (**Supplementary Figure 6**). Wind impact may be driven by the spillover of microbes from neighboring plants and air-driven transport of microbes ^[34]^, but also indirectly driven by the impact on neighboring plant communities, which may modify soil nutritional status ^[35, 36]^. Therefore, lower wind may be related to lower invasion of external microbes to the core microbiota and, with that, an increase in the relative abundance of the resident microbes of the core microbiota. Importantly, we observed both negative and positive correlations between ASVs belonging to core groups and environmental factors (**Figure 4b**). This may suggest that within core groups, there is an ASV balancing to adapt to specific environments and/or algal hosts. Whether this balancing dynamics is variable over time will require further investigation.

While ASV-level analysis allows us to explore environmental effects on within-core diversity across habitats, core members are systematically found together, and therefore community-level studies become more relevant. Network analysis is a powerful tool to explore microbe-microbe interactions across environments, rather than single-microbial groups independently. Here, network analysis has allowed us to demonstrate that not all core groups have the same behavior in a network context, and that there are probably different levels of interactions among core members and between core members with other members of the local microbiota (**Figure 5, Supplementary Figure 7**, **Supplementary Table 4**). This, together with the fact that only few of the core microbiota groups are significantly enriched in the presence of a photosynthetic host (**Supplementary Figure 2d, f**), could suggest that direct host selection may operate on only a fraction of the core microbiota. The remaining core members may instead be recruited indirectly through microbial interactions, metabolic dependencies, or community-level niche construction established by host-responsive taxa, consistent with models of hierarchical community assembly and priority effects ^[37]^.

We also observed that bacterial and fungal core groups have more negative interactions than non-core members, suggesting a potential negative feedback between core members of different kingdoms (**Figure 5a, b, Supplementary Figure 7a**). It had been previously observed that negative interactions predominate between bacteria and fungal members of the root microbiota ^[38, 39]^, where bacterial members, together with host metabolic inputs, regulate potential fungal microbiota dysbiosis ^[40]^. Modelling and experimental work has demonstrated that ecological competition stabilizes microbial interactions and that antagonistic interactions are prevalent in communities with niche overlap ^[41, 42, 43]^. Therefore, the fact that we observe stronger negative interactions within the core microbiota may reflect a stabilizing process in a microbial community that colonizes a common niche. Interestingly, these interactions may be environment-dependent, since we observed that different core orders are important for network structure among the three environmental clusters (**Figure 5c**, **d**). Further experimental validation will be necessary to explore the modularity and environment-dependent plasticity of the core microbiota in association with photosynthetic hosts.

## Conclusion

In this study, we characterized the natural diversity of the core microbiota associated with photosynthetic organisms by leveraging natural soil algal populations across diverse habitats. Our system reveals that although several bacterial and fungal orders consistently occur in association with photosynthetic hosts, their composition can vary substantially, at least in algal populations, across environments at finer taxonomic resolution. This variability arises from both shifts in ASV composition and changes in microbe–microbe interactions within core microbiota networks, highlighting the dynamic nature of the core microbiota in natural environments.

However, our study does not address the extent to which host identity contributes to shaping ASV-level core microbiota structure, whether different ecological interactions exist within core networks, or whether functional redundancy underlies the persistence of core microbial orders. Future work should therefore aim to disentangle these complex ecological and functional interactions to better understand the mechanisms maintaining core microbiota across photosynthetic organisms and environments.

## Methods

### Algal population sampling

One hundred forty-one natural sites across the southwest of France were selected to survey natural algal populations. Surface soil samples were taken into collection tubes as described in ^[6]^. Briefly, during the first two weeks of May 2023, soil patches where obvious green was visible, not belonging to moss patches, were selected. This was important for differentiating germinating mosses from true algal lawns. The soil surface was scratched using a metal spatula, which was rinsed with 70% ethanol between each site. Four independent samples were taken for each site and transferred to 1.2mL collection tubes in 96-well matrix racks. Samples were transported at room temperature during the day and stored at −80°C upon arrival at the laboratory until further processing.

For the comparison between surface and deeper soil layers, three sites out of the 141 were selected randomly. There, in addition to soil surface samples, the top 10cm layer of soil was removed, and three deeper soil samples were taken at each site.

### Environmental data and environmental clusters

The 141 natural sites selected had been described and characterized in previous work (e.g. in ^18, 19, 20^). We retrieved soil properties and plant community descriptors (i.e. diversity and composition) from the publicly available datasets. Additionally, we gathered meteorological data from weather stations near the study sites, from one, two, three, and four weeks before our sampling dates (**Supplementary Figure 1, Supplementary Table 1**).

Given the variability of environmental conditions across natural sites, we grouped them into environmental clusters (ECs). For this, we selected all sites for which we had the full datasets (100/141). Sites were therefore clustered based on meteorological conditions, soil physicochemical properties, and plant community descriptors. Environmental data were imported into *R* and separated into three predefined variable groups: (i) meteorological variables (dew point, humidity, precipitation, pressure, temperature, wind metrics), (ii) soil variables (nutrients, elemental composition, pH, water holding capacity, organic matter), and (iii) plant descriptors extracted from ^[20]^. The first PCoA axis of plant communities is associated with (i) annual species occurring in bare tilled, fallow or recently abandoned arable lands, and (ii) perennial species occurring in mesic grasslands ^[20]^. The three datasets were combined and analyzed using Multiple Factor Analysis (MFA) implemented in the *FactoMineR* package, with each data block treated as a separate group to balance their contributions. The optimal number of clusters was assessed using the elbow method based on the total within-cluster sum of squares (WSS) across *k* = 1–15. Initial inspection suggested four ECs; however, cluster 4 had insufficient sample representation and clustering was recomputed, resulting in a final solution of three ECs (**Supplementary Figure 1**).

### DNA extraction and amplicon sequencing library preparation

DNA was extracted using a modified version of a homemade protocol ^[44]^ for 1.2mL collection tubes. Briefly, cells were lysed using a pre-warmed buffer at 65°C, which contains 100mM Tris-HCl, 100mM NaCl, 10mM EDTA pre-mix, to which 1.5% SDS, 40mM DTT, and 100µg/mL proteinase K was added right before utilization. 300µL of the lysis buffer were added to each sample, which were then ground in a FastPrep-90 (MP Biomedicals) for 30sec at 6 m/s. Samples were then incubated at 65°C for one hour. Before spinning down the samples for 5min at 1000RCF, they were mixed by inversion 20 times. Then, 125µL of the supernatant was transferred to a new 96-deep-well plate, and 46.5µL of potassium acetate 5M was added. Samples were then centrifuged at 6200RCF for 10min, and the supernatant (∼110µL) was then transferred to a new 96-deep-well, and mixed with SeraMag SpeedBeads® Carboxyl Magnetic Beads (GE Healthcare; #65152105050450) for purification. Samples were mixed with beads by inversion 20 times, and then placed on a magnet. After the beads were attached to the magnet, the supernatant was removed, followed by two consecutives washes of ethanol 80%. Then, samples were then eluted in 50µL of sterile MiliQ water.

Amplicon sequencing libraries were built by following a two-step amplification protocol. In the first step, target-specific primers were used, to which internal tags and Illumina adaptors were added to allow further multiplexing of the samples (**Supplementary Table 3**). 16S, ITS2, and 18S target genes were used to describe bacterial, fungal, and eurkaryotic communities, respectively. Each sample was amplified in a 10 μL reaction volume containing 0.1µL of GoTaq polymerase, 2.5µL of 5x GoTaq buffer (Promega), 0.1µL of 50mg/mL BSA, 0.25µL of 10µM dNTPs (Promega), and 0.25µL of 10μM forward and reverse primers. PCR was performed using the following parameters: 94°C/2 min, 94°C/30 s, 59°C/30 s, 72°C/60 s, 72°C/5 min for 30 cycles. Afterwards, single-stranded DNA and proteins were digested by adding 0.05 μL of Exonuclease I and 0.1µL of Shrimp Alkaline Phosphatase (New England BioLabs), mixed with 3.85 μL of sterile MilliQ water to the PCR product. Samples were incubated at 37°C for 30 min and enzymes were deactivated at 85°C for 15 min. 5μL of this reaction were used for a second PCR, where each sample was amplified in a 15μL reaction volume containing 0.125µL of GoTaq polymerase, 2.5µL of 5x GoTaq buffer (Promega), 0.25µL of 10µM dNTPs (Promega) and 0.5µL of 10μM P5 Illumina TruSeq forward primer. 0.5µL of indexed reverse Illumina TruSeq primers were individually added to each reaction. PCR was performed using the following parameters: 94°C/2 min, 94°C/30 s, 59°C/30 s, 72°C/60 s, 72°C/5 min for 10 cycles. PCR samples were then pooled by mixing 7.5µL of each reaction per plate and quality was controlled by loading 5μL of each pool of samples on a 2% agarose gel and confirming that no band was detected for negative controls. Then, 80μL of the pooled library was loaded in a 2% agarose gel and run for 2h at 50V. Subsequently, bands with a size of ∼300-500 bp were cut out and purified using the Wizard® SV Gel and PCR Clean-Up System (Promega). The final library concentration was estimated fluorescently (Qubit Fluorometer, ThermoFisher Scientific). Paired-end Illumina sequencing was outsourced to Novogene.

### Analysis of amplicon sequencing data

Amplicon sequencing data were demultiplexed according to their barcode sequence added in the second PCR by the sequencing company. Upon data delivery, sequencing data were further demultiplexed by the internal tags utilized in the first PCR using *cutadapt* (v4.6). Afterwards, QIIME (version 2021.11, ^[45]^) was used to process the raw sequencing reads of each sample. Unique amplicon sequencing variants (ASVs) were inferred from error-corrected reads, followed by chimera filtering, using the DADA2 pipeline (v 1.30.0). Next, ASVs were aligned to the SILVA database for taxonomic assignment of bacterial and eukaryotic reads (version 138, ^[46]^), and the UNITE database for a taxonomic assignment of fungal reads (version 8, ^[47]^), using the naïve Bayesian classifier implemented by DADA2. Raw reads were mapped to the inferred ASVs to generate an abundance table, which was subsequently employed for analyses of diversity and differential abundance under the *R* environment (v4.5.2).

### Compilation of public datasets

Based on the methodological framework described above, this approach was applied to multiple datasets derived from published studies on photosynthetic organisms (**Supplementary Table 2**), as well as to our data from terrestrial algal communities. Each dataset was processed using the same preprocessing strategy to ensure comparability among studies. ASV abundance tables were normalized to relative abundance, and samples with insufficient sequencing depth were removed before rarefaction to an even sequencing depth of at least 2,000 reads, which was sufficient to retain the potential microbial diversity (**Supplementary Figure 8**). Following preprocessing, microbial profiles were integrated with the corresponding metadata to account for the sampling structure, and abundances were summarized at the site level. Taxonomic information was then incorporated to associate ASVs with their classifications, and microbial abundances were subsequently aggregated at the taxonomic rank of “order”. This produced comparable community profiles across datasets, which were used for downstream analyses.

### Abundance-occupancy calculation

To identify core members of microbial communities, we applied the abundance–occupancy framework described in ^[23]^. This method is based on the occupancy (proportion of samples in which a taxon is detected) and the relative abundance of each ASV. Taxa were ranked according to a combined index incorporating detection frequency and replication consistency. To determine the set of core members, Bray-Curtis similarity was calculated iteratively for all ranked taxa, with each taxon added one by one according to their ranking to assess its contribution to the overall dissimilarity of the community. The number of iterations depends on the total number of taxa in the dataset. As described in ^[23]^, two thresholds were used to define the core: (i) the elbow method, based on the first-order difference in similarity gain, and (ii) the last 2% increase in Bray-Curtis similarity.

### Network analysis

Networks were constructed from 16S and ITS amplicon sequencing data. To improve statistical robustness, a bootstrapping approach with replacement was applied, using random subsampling of samples or columns depending on the analysis. After each bootstrap, ASVs were matched to their full taxonomic annotation from a reference file including six levels: Kingdom, Phylum, Class, Order, Family, and Genus. A unique taxonomy string was created for each ASV and used as a row identifier in the abundance matrix, ensuring a clear link between ASVs and their biological identity. ASVs were filtered to keep only those present in more than 5% of samples and with a mean relative abundance above 0.00001. The filtered abundance matrix was re-normalized, and pairwise correlations were calculated using the Spearman method. This produced a matrix of correlation coefficients (*ρ*) and a matrix of associated *p-*values. Only correlations with *ρ* ≥ 0.2 and *P* < 0.01 were kept, and self-correlations were removed. To create the network, the filtered correlation and p-value matrices were converted into an edge table. Each row represented a significant link between two ASVs. Duplicate edges were removed by discarding the upper triangle of the correlation matrix, and only significant non-zero correlations (*P* < 0.01) were retained. The resulting table included the two connected ASVs and their correlation strength (edge weight). A separate node table was also generated, listing all ASVs involved in at least one edge along with their full taxonomy (Kingdom to Genus). Node attributes were extracted from Cytoscape (version 3.10.1) for visualization and analysis.

Network topological metrics were normalized before visualization to allow comparison across properties with different scales. Betweenness centrality, degree, closeness centrality, average shortest path length, clustering coefficient, eccentricity, neighborhood connectivity, and topological coefficient were rescaled using min–max normalization to a range of 0–1. Significant differences between metrics were calculated either between core *vs* non-core nodes or between core order nodes, across ECs networks following Kruskal-Wallis and pairwise *post-hoc* Wilcoxon tests (BH-corrected), using the non-scaled values.

### Network comparison and order-specific permutation

Networks were inferred for each environmental cluster (EC). To ensure robustness and reduce the bias introduced by sample number, a subset of 15 samples was randomly selected from each EC, and EC-specific networks were constructed using 11 bootstrap replicates for each pairwise comparison. For each bootstrap iteration, Spearman rank correlations were computed between microbial taxa, and the resulting correlation matrix was used as the network adjacency matrix. The eigenvectors of each adjacency matrix were then computed and used for calculating pairwise Euclidean distances between networks across ECs to quantify network differences. Principal component analysis (PCA) was then applied to visualize the multidimensional network distances in a reduced space, enabling the assessment of network divergence across environmental conditions. As network distances were calculated separately for each pairwise EC comparison (e.g. EC1 *vs* EC1, EC1 *vs* EC2, EC1 *vs* EC3), with 11 bootstrap replicates performed for each comparison; therefore consequently, each EC contributed to four pairwise comparisons and generated 44 inferred networks (4 comparisons × 11 bootstrap replicates). All inferred networks were included in the PCA based on eigenvector distances, resulting in 44 observations for each EC in **Figure 5c**.

To identify the taxonomic orders most responsible for the observed network differentiation, we employed an order-specific permutation test. In each permutation, while sample labels for a given taxonomic order were randomly shuffled, all other taxa remained unchanged, thereby disrupting the association patterns of that order without altering the overall community structure. The network distances between ECs were recalculated for each permuted network and compared to those derived from the original (non-permuted) networks. The contribution of each order to network divergence was quantified by the changes in pairwise network distances induced by permutation. Orders were ranked according to their average effect size across permutations, with higher values indicating greater influence on the structural separation of EC-specific networks.

## Data availability

All processed data and code used to produce the manuscript’s figures are available at https://github.com/duranpa/EcoDrivers_Coremicrobiota_Garriguesetal. In addition, all raw reads are deposited and available at the ENA, under the accession PRJEB108334.

Generative AI (ChatGPT, OpenAI) was used as an assistance tool for refining, and troubleshooting R scripts. The authors independently reviewed the generated code and verified all analytical procedures and results.

## Author contribution and funding

P.D and F.R were involved in conceptualization. P.D and R.G. conceived and designed the analysis. P.D., F.R., I.L. and E.W. performed the harvesting survey. P.D. and I.L. performed DNA extractions and sequencing libraries preparation. V.G., R.G., N.P. and P.D. performed data curation and formal analysis. The manuscript was written by P.D., with input from all authors. The final manuscript was approved by all authors.

This project has received funding from the European Research Council (ERC) under the European Union’s Horizon 2020 research and innovation program (grant agreement No 951444 – PATHOCOM). This study was performed at the LIPME belonging to the Laboratoire d’Excellence (LABEX) entitled TULIP (ANR-10-LABX-41). R.G. was supported by the European Research Council H2020 StG (erc-stg-948219, EPYC) and BBSRC Institute Strategic Programme Food Microbiome and Health (BB/ X011054/1, BBS/E/F/000PR13631).

## Acknowledgments

We would like to thank Dr. Léa Frachon for providing the plant cover measurements across the surveyed sites. We would also like to thank Dr. José Flores-Uribe for useful discussions and suggestions with data analysis, as well as for critical reading of this manuscript.

## Supplementary figures

**Supplementary Figure 1:**
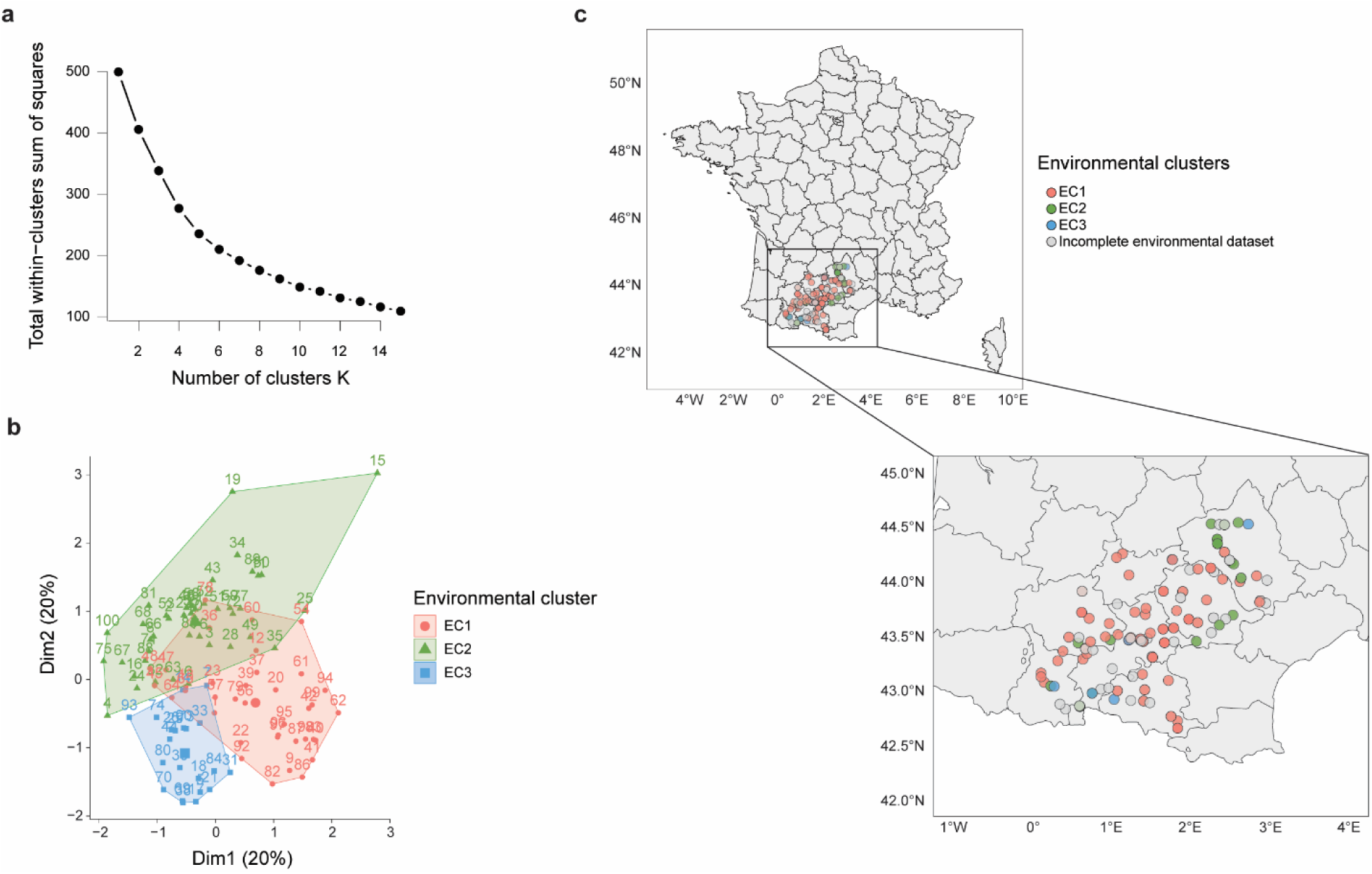
Surveyed sites across the Southwest of France. The 100 sites for which we have a full dataset for environmental factors, can be grouped by environmental clusters. **a)** Elbow method to calculate how many clusters best explain the differences between sites. **b)** Principal Components Analysis of the 100 sites, clustered based on the environmental composition (soil properties, climatic conditions and plant communities). **c)** Map of the 141 sites used in this study. 41 of these sites do not have complete datasets, either on soil properties, plant communities’ composition or climatic conditions and therefore were removed from the environmental clustering (grey points).

**Supplementary Figure 2:**
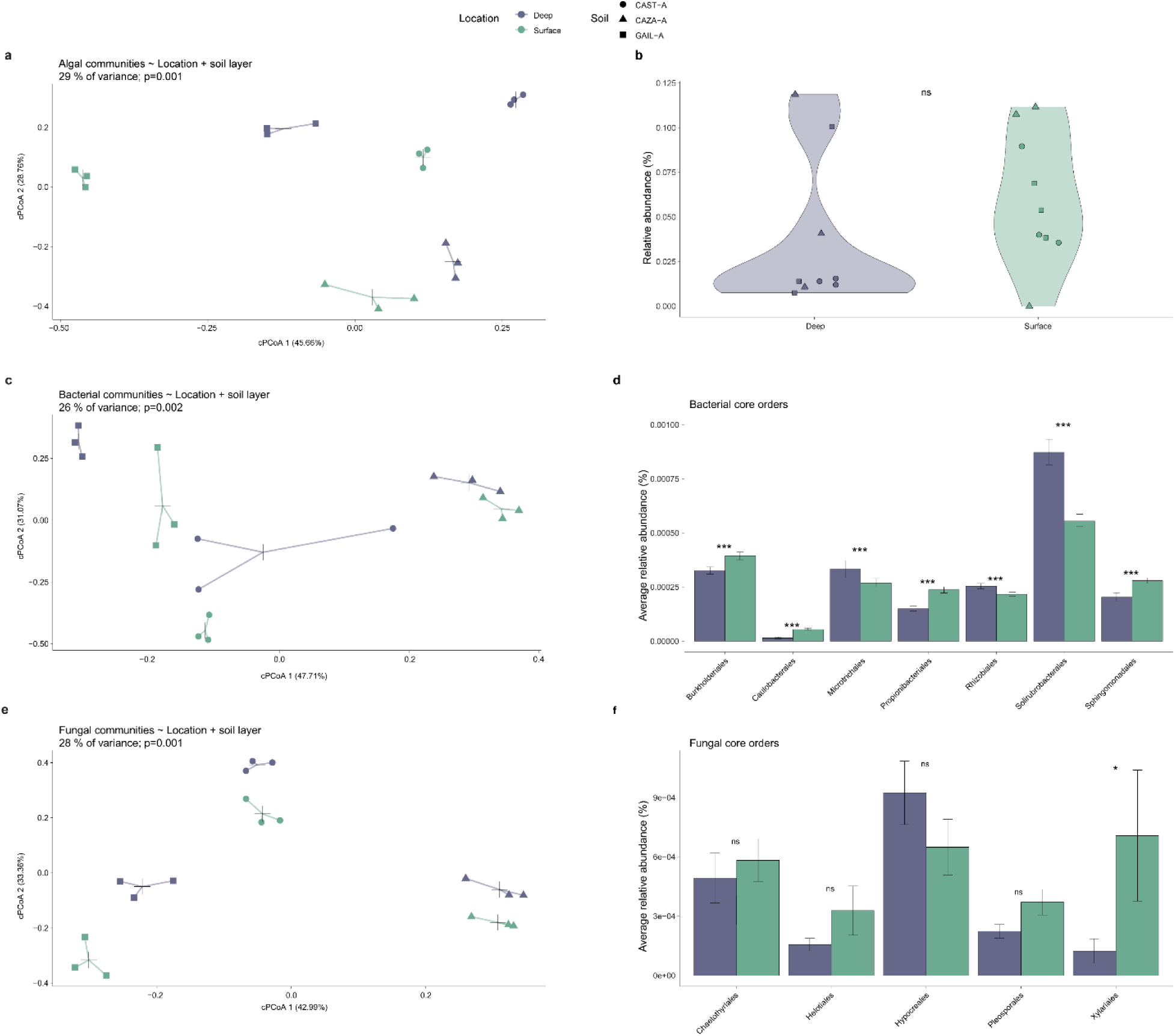
Algal and microbial communities on deep soil layers *vs* photosynthetic surface layer. **a)** Principal Component Analysis of algal community based on Bray-Curtis distances, constrained by soil layer (“Deep” and “Surface”) and harvesting site (PERMANOVA test, *P* = 0.001). **b)** Relative abundance of green algae across “Deep” and “Surface” soil layers, with data points categorized by soil type (CAST-A, CAZA-A and GAIL-A) (n.s., *P* > 0.05 Wilcoxon Test, Holm-corrected) **c)** Principal Component Analysis of bacterial community based on Bray-Curtis distances, constrained by soil layer (“Deep” and “Surface”) and harvesting site (PERMANOVA test, *P* = 0.001). **d)** Relative abundance of bacterial core orders between Deep and Surface soil layers. Significant differences are depicted with asterisks (***, *P* < 0.001 Wilcoxon Test, Holm-corrected). **e)** Principal Component Analysis of fungal community based on Bray-Curtis distances, constrained by soil layer (“Deep” and “Surface”) and harvesting site (PERMANOVA test, *P* = 0.001). **f)** Relative abundance of fungal core orders between Deep and Surface soil layers, showing stable distribution for most orders (ns, *P* > 0.05 Wilcoxon Test, Holm-corrected), except for Xylariales (*, *P* < 0.05 Wilcoxon Test, Holm-corrected).

**Supplementary Figure 3:**
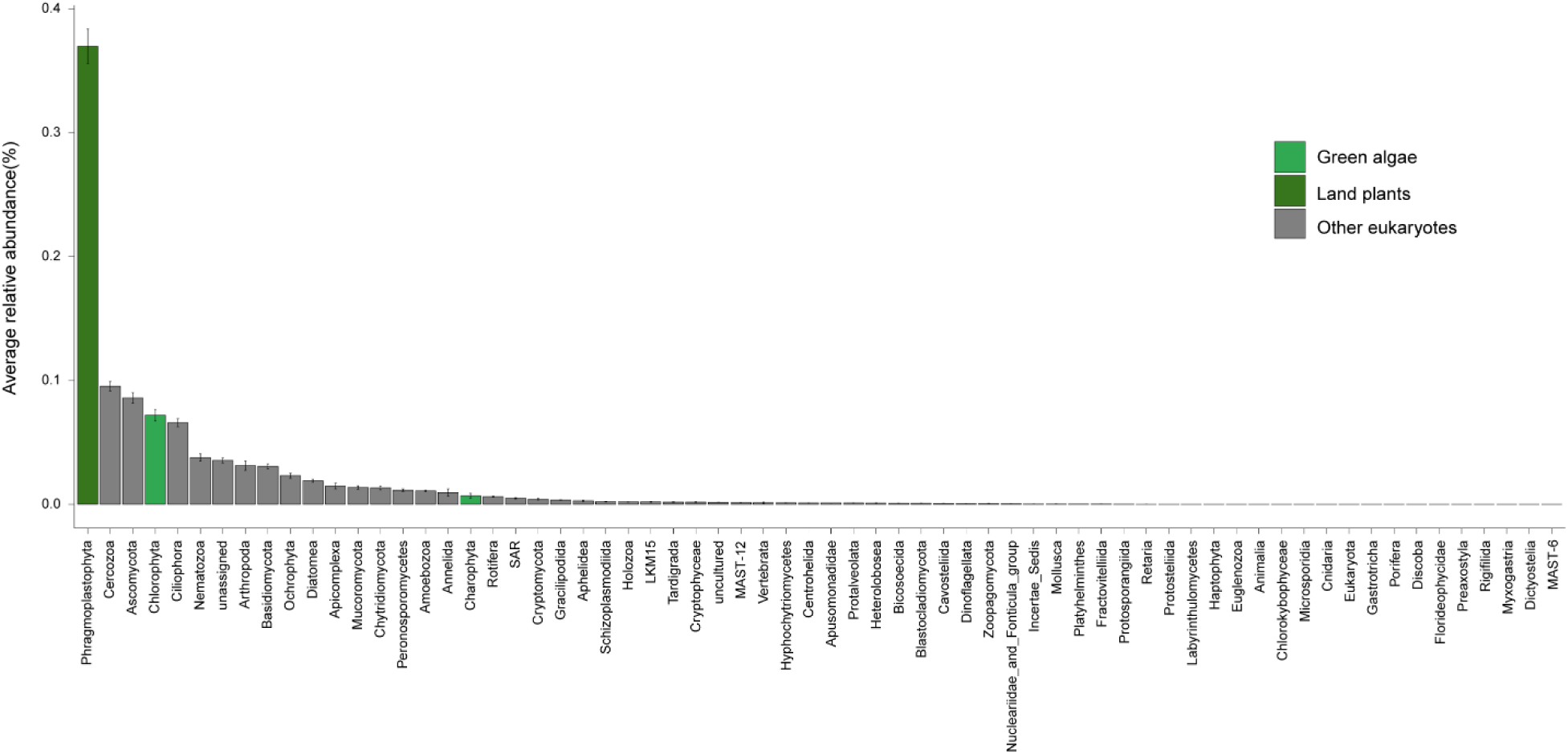
**a)** Average relative abundances of eukaryotic phyla, identified using the 18S target gene. Land plants and green algae are highlighted in green. Mean values across all samples (n=564) are depicted here, with whiskers indicating standard error.

**Supplementary Figure 4:**
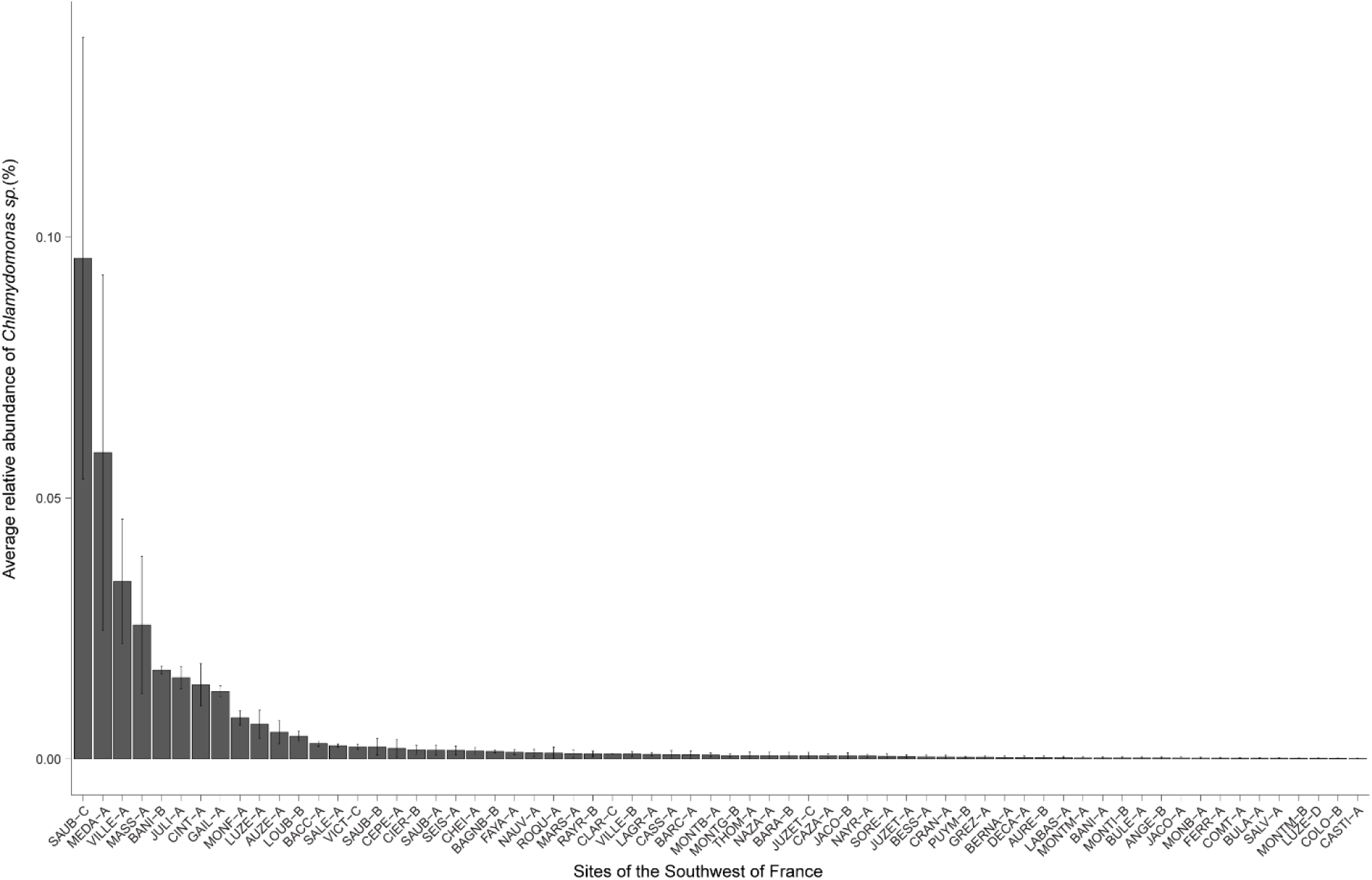
Members of the *Chlamydomonas* genus are found across 66 of the surveyed sites. In the figure, the average relative abundance of the *Chlamydomonas* genus is depicted in each site (n=4), with whiskers indicating standard error.

**Supplementary Figure 5:**
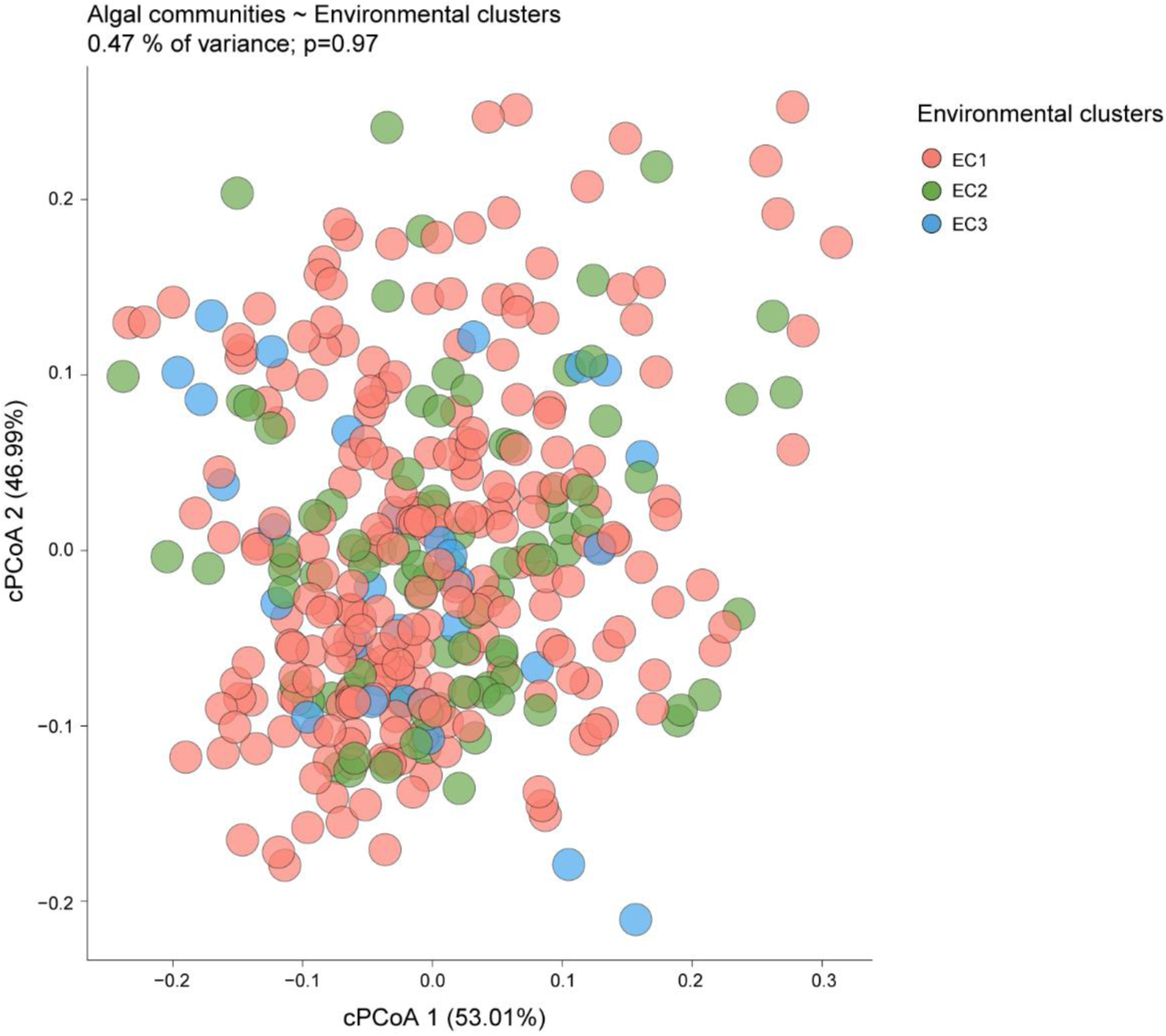
Environmental clusters do not drive algal communities’ structure. Principal Components Analysis of algal communities, constrained by the environmental clusters to which each natural site belongs to. A permutational analysis of the variance (PERMANOVA) showed a non-significant effect of the environmental cluster on algal community composition (*P* = 0.97).

**Supplementary Figure 6:**
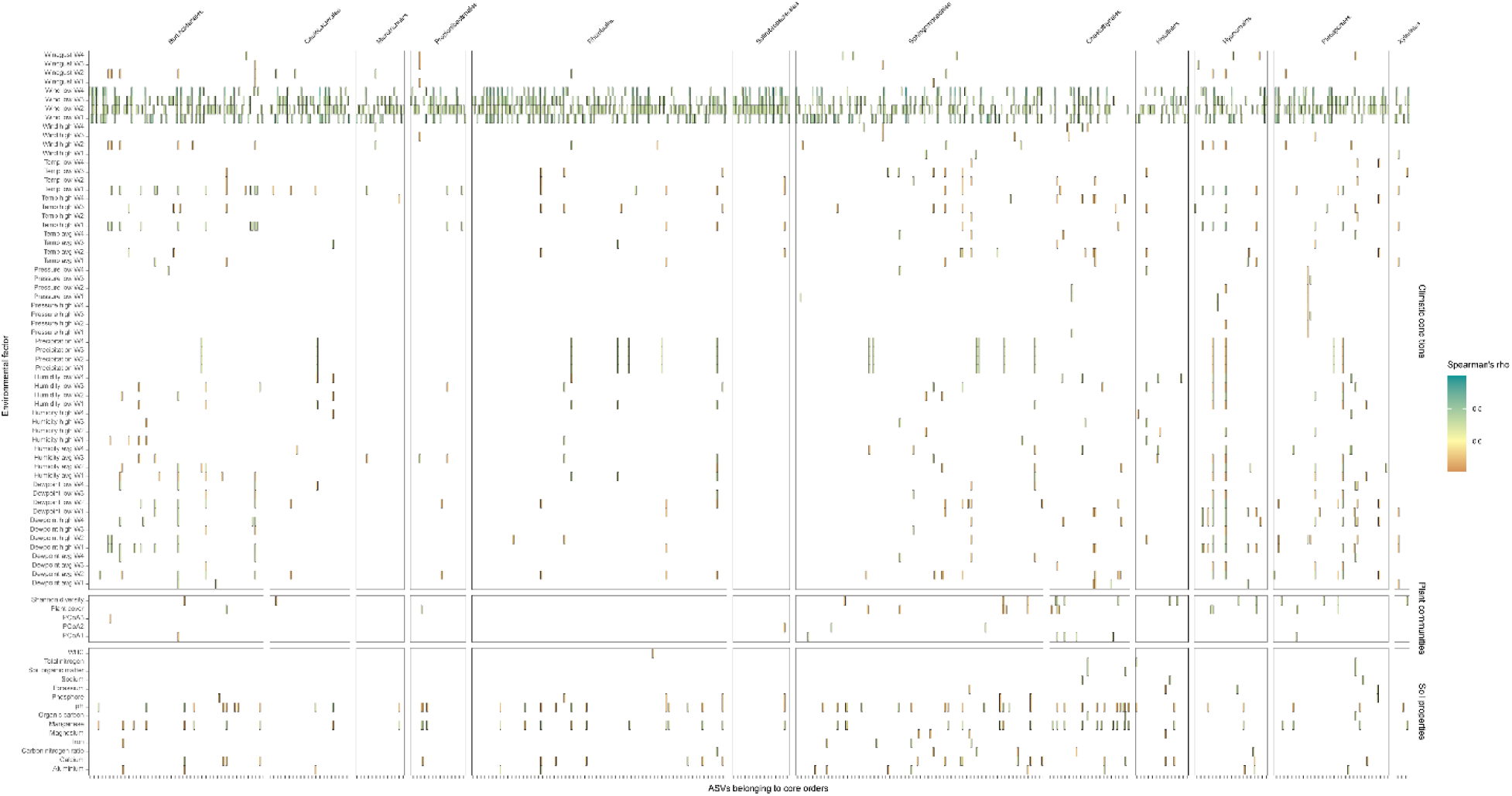
Correlation between environmental factors and the relative abundance of ASVs belonging to core orders. Only significant correlations are shown here (Spearman’s correlation, *P* < 0.05, FDR-corrected), color-coded based on the strength of the correlation.

**Supplementary Figure 7:**
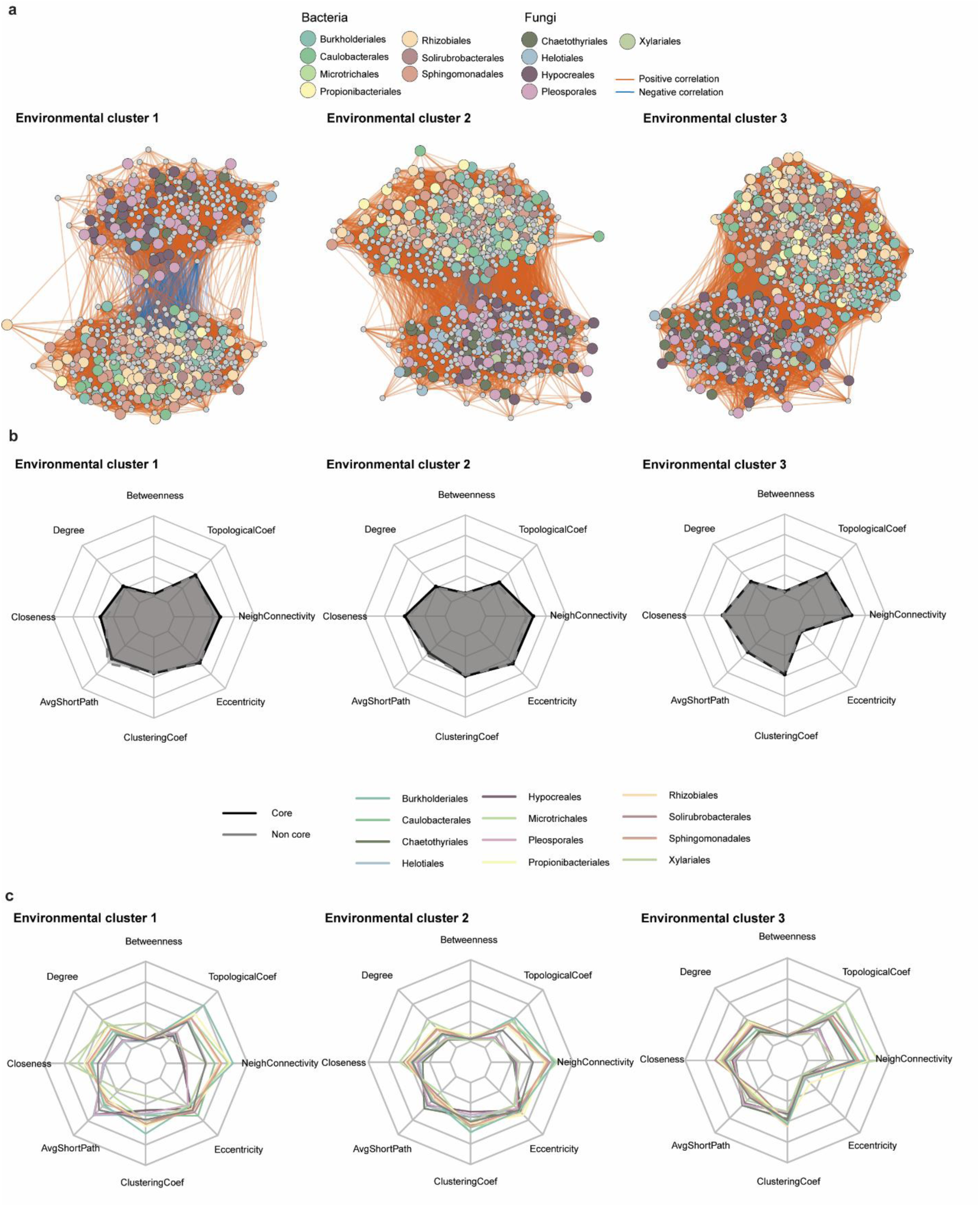
The core microbiota forms a complex network of interactions that varies across environmental clusters (ECs). **a)** Network of interactions across natural sites for the three environmental clusters. The leftmost network is the same as in Figure 5a. Colors in nodes indicate whether a given node belongs to a core order (colored), or any other order (grey). Negative correlations are depicted with blue edges, and positive correlations are depicted with orange edges. Only significant correlations are included here (Spearman correlation, *P* < 0.01) **b)** Network properties of core *versus* non-core nodes across the three environmental clusters (ECs). **c)** Network properties of each core order *versus* non-core nodes across the three environmental clusters (ECs). Network topological metrics were normalized prior to visualization to allow comparison across properties with different scales.

**Supplementary Figure 8:**
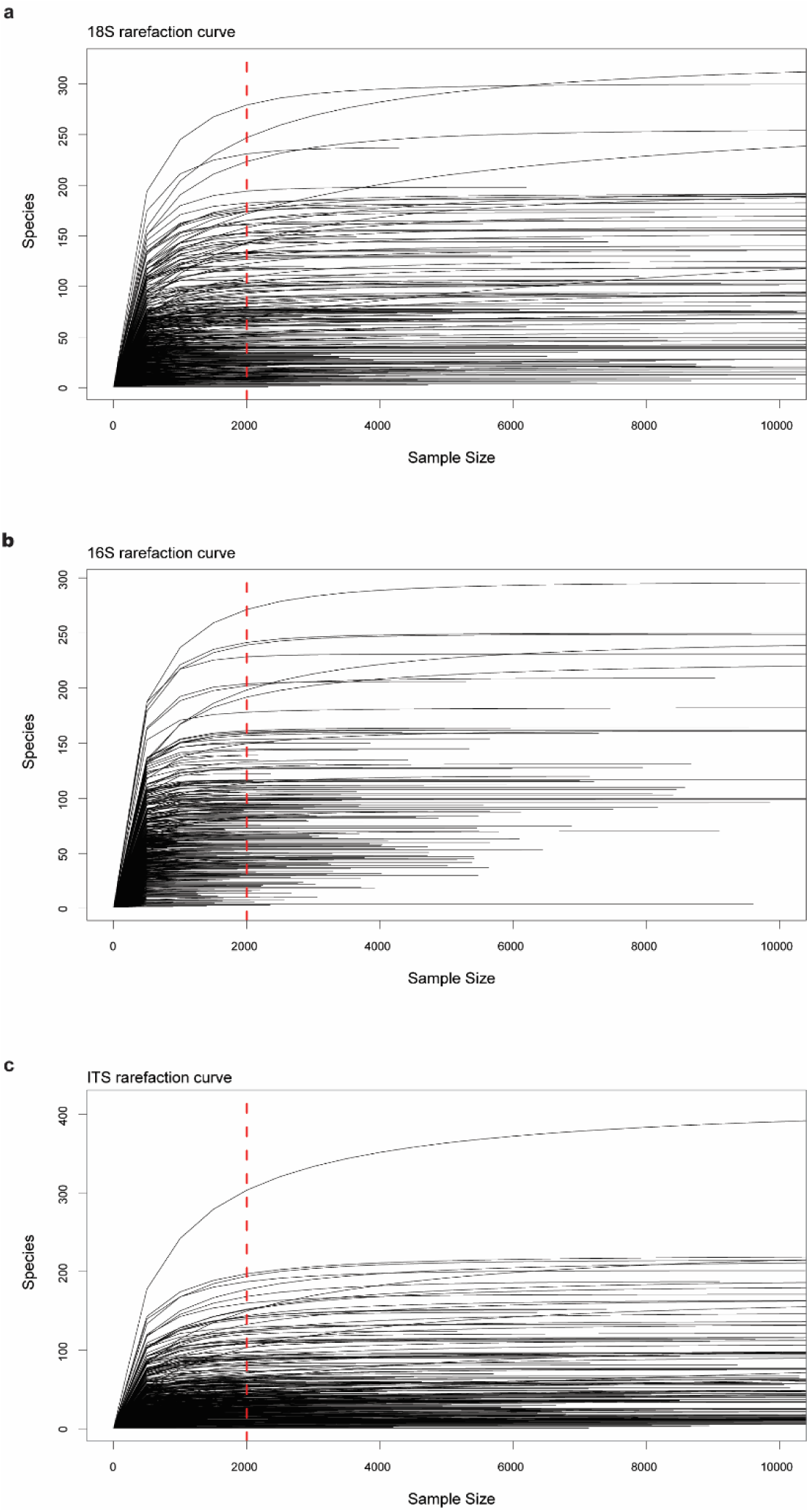
Rarefaction curves of amplicon sequencing dataset, for 18S (a), 16S (b) and for ITS (c). The dashed red line indicates the minimum depth (2000 reads) that each sample needed to have to be retained for further analysis. In all datasets, 2000 reads were higher than the minimum number of reads to retain the potential microbial diversity (elbow method).

## Supplementary tables

**Supplementary Table 1:** Environmental factors from sites across the Southwest of France, including soil properties, climatic factors one, two, three and four weeks prior to harvesting, and plant communities’ descriptors.

**Supplementary Table 2:** Datasets to define the core microbiota of photosynthetic organisms

**Supplementary Table 3:** Internally-tagged primers used in PCR1 for multiplexing

**Supplementary Table 4:** Kruskal-Wallis and pairwise *post-hoc* Wilcoxon test (BH-corrected) across network topologies, for core vs non-core nodes across environmental clusters and per core order.

